# Polymerase ζ activity is linked to replication timing in humans: evidence from mutational signatures

**DOI:** 10.1101/011494

**Authors:** Vladimir B. Seplyarskiy, Georgii A. Bazykin, Ruslan A. Soldatov

**Author notes:** These authors contributed equally to this paper (Vladimir B. Seplyarskiy, Ruslan A. Soldatov).

## Abstract

Replication timing is an important determinant of germline mutation patterns, with a higher rate of point mutations in late replicating regions. Mechanisms underlying this association remain elusive. One of the suggested explanations is the activity of error-prone DNA polymerases in late-replicating regions. Polymerase ζ (pol ζ), an essential error-prone polymerase biased towards transversions, also has a tendency to produce dinucleotide mutations (DNMs), complex mutational events that simultaneously affect two adjacent nucleotides. Experimental studies have shown that pol ζ is strongly biased towards GC->AA/TT DNMs. Using primate divergence data, we show that the GC->AA/TT pol ζ mutational signature is the most frequent among DNMs, and its rate exceeds the mean rate of other DNM types by a factor of ~10. Unlike the overall rate of DNMs, the pol ζ signature drastically increases with the replication time in the human genome. Finally, the pol ζ signature is enriched in transcribed regions, and there is a strong prevalence of GC->TT over GC->AA DNMs on the non-template strand, indicating association with transcription. A recurrently occurring GC->TT DNM in HRAS gene causes the Costello syndrome; we find a 2-fold increase in the mutation rate, and a 2-fold decrease in the transition/transversion ratio, at distances of up to 1 kb from the DNM, suggesting a link between the Costello syndrome and pol ζ activity. This study uncovers the genomic preferences of pol ζ, shedding light on a novel cause of mutational heterogeneity along the genome.

## Introduction

Germline mutations are the source of material for the evolutionary processes and the cause of genetic diseases. The mutation rate per generation is determined by an interplay between several factors: number of cell divisions, extracellular and intracellular DNA damage, accuracy of DNA-synthesizing polymerases, and DNA repair fidelity. Furthermore, the human mutation rate varies along the genome (reviewed in Hodgkinson and Eyre-Walker 2011; Ségurel et al. 2014); the mechanisms underlying this variation are only partially understood. Factors correlated with the local mutation rate include replication timing, meiotic recombination rate, GC content, DNA accessibility, and distance from the telomeres. Human DNA replication is a complex process which involves synthesis of DNA from multiple origins, so that replication of different large-scale regions is activated in a robust temporal order which is conserved between tissues and species (Ryba et al. 2010; Yaffe et al. 2010). Replication timing is among the strongest determinants of the mutation rate, as reflected by the evolutionary divergence of primates (Stamatoyannopoulos et al. 2009; Chen et al. 2010) or insects (Weber et al. 2012) and the somatic mutational patterns in humans (*e.g.* Koren et al. 2012; Woo and Li 2012). Replication timing affects multiple classes of mutations, including all kinds of point mutations (Chen et al. 2010) and copy number variation induced by nonallelic homologous recombination or nonhomologous end-joining (Koren et al. 2012).

Here, we focus on multinucleotide mutations (MNMs). MNM is a mutation event that simultaneously changes several nucleotides in a row. Mechanisms and factors governing the distribution of these events along the genome are unknown. The most frequent class of MNMs is dinucleotide mutations (DNMs) (Schrider et al. 2011; Terekhanova et al. 2013) which affect two sites at once. The rates of DNMs were estimated from de-novo mutations in parent-child trios, interspecies divergence, intraspecies polymorphism and disease causing mutations (Table 1 in Chen et al. 2014). The obtained estimates were rather similar, placing the rate of DNMs at about 0.4% of the single nucleotide mutation (SNM) rate. The GC->AA/TT mutation is the most frequent DNM in human polymorphism (Harris and Nielsen 2014) and disease data (Chen et al. 2013). The same DNM is strongly associated with the activity of polymerase ζ (pol ζ) (Stone et al. 2012).

Pol ζ is able to perform translesion synthesis (TLS) and to extend the non-perfectly matched primers (Lawrence and Maher 2001). It is recruited to bypass replication stalling associated with double strand breaks (Sharma et al. 2012), mismatches (Northam et al. 2010) or DNA non-B structures (Northam et al. 2014). Pol ζ is the only TLS polymerase that is essential for embryonic development in mammals (Bemark et al. 2000; Lange et al. 2012). The mutation rate of pol ζ is about 10^−3^ point mutations per site, making it substantially more error-prone than other homologous B family polymerases α, δ and ε (Zhong et al. 2006; Sakamoto et al. 2007) which play the central role in DNA replication (Waisertreiger et al. 2012). Moreover, the mutational spectrum of pol ζ is enriched in transversions (Stone et al. 2012).

Experimental studies in yeast (Harfe and Jinks-Robertson 2000; Stone et al. 2012; Northam et al. 2014) and mammals (Saribasak et al. 2012) suggest that pol ζ tends to produce DNMs. Nucleotide excision repair (NER) deficient *S. cerevisiae* line with error-prone pol ζ demonstrates a strong overrepresentation of complementary GC->AA and GC->TT DNMs, and also of TC->AA, among the detected DNMs (Stone et al. 2012). However, the overrepresentation of TC->AA is probably a result of an ascertainment bias due to these mutations being more likely to be detected in assays. Indeed, the rate of TC->AA strongly exceeds the rate of its GA->TT complement (Stone et al. 2012); this and other considerations suggest that the same bias holds for the mutations observed in human disease (Chen et al. 2013). Mammalian experimental studies also suggest that the WGCW context (where W denotes A or T) is a hotspot of tandem mutations associated with pol ζ (Saribasak et al. 2012). Therefore, the GC->AA/TT DNM seems to represent the mutational signature of pol ζ (Harris and Nielsen 2014). This allows us to use the rate of GC->AA/TT DNMs to trace the activity of pol ζ along the human genome.

In this study, we show that the GC->AA/TT is the most common DNM in human-chimp divergence. The pol ζ signature is especially pronounced in introns, where a strong excess of GC->TT over GC->AA on the non-template strand is observed, suggesting association with transcription. The pol ζ signature is also stronger at hypermutable CpG sites, controlling for the confounding factors. While the rate of all DNMs is only very weakly associated with replication timing, the pol ζ signature rate is radically higher in late replicating compared to early replicating regions. Together, these finding indicate the heterogeneity of the genomic effect of pol ζ, shedding light on a novel cause of mutational heterogeneity along the genome.

## Results

As a proxy for the human mutation rate, we used the level of human-chimpanzee divergence. Using orangutan as the outgroup allows us to distinguish between mutations that occurred in the human and in the chimpanzee lineage. We limited our analyses to the putatively neutrally evolving sites by excluding exons, 5‵ UTRs, 3‵ UTRs and 50 bp intron flanks. In the main analysis, we also masked the CpG sites which are hypermutable due to extensive cytosine methylation (Bird 1980), whereas in the analysis of MNMs, we excluded the tandem substitutions that may occur through the CpG intermediate state (see Methods). Using 50 kb non-overlapping windows along the genome, we estimated mutation rate, GC content, replication timing, DNA accessibility, H3K9me3 and meiotic recombination rate.

### Dinucleotide mutations

Two substitutions at adjacent sites in the human lineage after the last common ancestor with chimpanzee could arise either due to a DNM or to two independent SNMs. Conversely, substitutions at adjacent sites such that the first substitution occurred in the chimpanzee lineage, and the second, in the human lineage, could only arise as independent SNMs (Fig. 1A). This allows us to determine the fraction of DNMs among all observed human-chimpanzee tandem differences (Terekhanova et al. 2013; see Methods). We observed 41,234 differences at adjacent sites with both mutations at the human branch (tandem same branch mutations, TSBMs), and 22,905 differences at adjacent sites with mutations at different branches (tandem different branch mutations, TDBMs) (Fig.1A). In the absence of DNMs, the number of TDBMs should be similar to the number of TSBMs, since the lengths of the two branches are nearly equal (Chimpanzee Sequencing and Analysis Consortium 2005); DNMs should lead to an increase in the number of TSBMs but not of TDBMs. Therefore, almost half of the TSBMs occurred as DNMs (Fig. 1B). The estimated DNM rate is 0.41% of the SNM rate, in line with our previous results (Terekhanova et al. 2013) as well as with those of others (reviewed in Chen et al. 2014).

**Figure 1.**
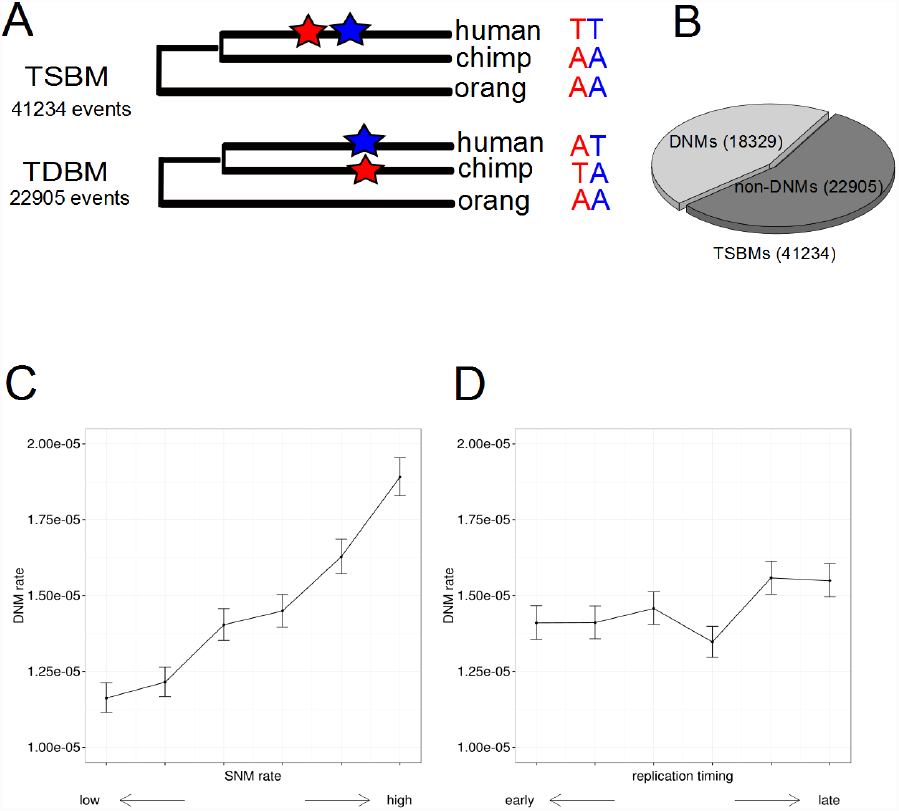
Properties of DNMs. (A) Schematic representation of divergence of human and chimpanzee from their last common ancestor. TSBMs are mutations that occurred at the same (human) phylogenetic branch; TDBMs are mutations that occurred at different branches. The number of DNMs can be estimated as the excess of TSBMs over TDBMs. (B) DNMs comprise a substantial part (44%) of TSBMs. (C) DNM rate is strongly associated with SNM rate (p<2 *10^−23^). (D) DNM rate is only weakly associated with RT (p=0.03).

Human-chimp divergence is a reliable source of information on the rate of DNMs. First, it carries orders of magnitude more DNMs than can be obtained from data on *de novo* mutations inferred from parent-offspring comparisons (a total of 7 MNMs at distance not exceeding 20 nucleotides, Schrider et al. 2011) and disease-associated variants (a total of 448 DNMs, Chen et al. 2013). Second, the phylogenetic framework is a simple yet reliable approach, in contrast to complicated models required to obtain DNM rates from human nucleotide diversity (Harris and Nielsen 2014). Third, our approach uses high quality reference genomes. Finally, non-coding substitutions in the human lineage are only slightly affected by selection (Rands et al. 2014), making our results less biased, compared to estimations from human disease-causing mutations and even polymorphism data where selection affects neutral sites due to linkage (Charlesworth 2009).

As Harris and Nielsen (2014), we observed that DNMs are enriched in transversions, compared to SNMs; specifically, the transition/transversion ratio is 60% lower among TSBMs compared to TDBMs (χ^2^ test, p<1.1 *10^−67^). This contrast suggests the difference of mutational mechanisms underlying DNMs and SNMs. However, TDBMs are also enriched in transversions by 18% compared to non-tandem mutations (χ^2^ test, p<1.2 *10^−34^), in line with our previous study (Seplyarskiy et al. 2012).

Genomic regions with elevated SNM rates are substantially more prone to DNMs (Fig. 1C, p=2*10^−23^, Cochran-Armitage test for trend in proportions with permutation-based null distribution (permuted CA test). Therefore, both the DNM rate and the SNM rate co-vary on a 50Kb scale, implying that although the mechanisms causing them may be distinct, they are correlated. These and all subsequent results were independent on the number of bins chosen (Suppl. Table S1), indicating that they are not an artifact of the binning procedure. In contrast, the DNM rate is only weakly associated with late RT (Fig 1D, p<0.03, permuted CA test; Suppl. Fig S1). This weak association is, however, robust; in particular, it is not the consequence of differences in GC content of early *vs.* late replicating regions (Suppl. Table S1).

### GC->AA/TT signature of pol ζ

Experimental studies in yeast have shown that the GC->AA/TT DNMs are strongly overrepresented among the DNMs; this has been attributed to the activity of pol ζ (Stone et al 2012). Harris and Nielsen (2014) observed that GC->AA/TT is also the most common type of tandem mutations occurring at adjacent sites in the human polymorphism data (The 1000 Genomes Project Consortium 2010) in perfect LD. We observed that the GC->AA/TT DNMs are much more frequent in the human-chimp divergence than other types of DNMs (Fig. 2): among the GC->AA/TT TSBMs, DNMs comprise 79%, while among all TSBMs, DNMs comprise only 44% (χ^2^ test, p<1.7 *10^−68^).

**Figure 2.**
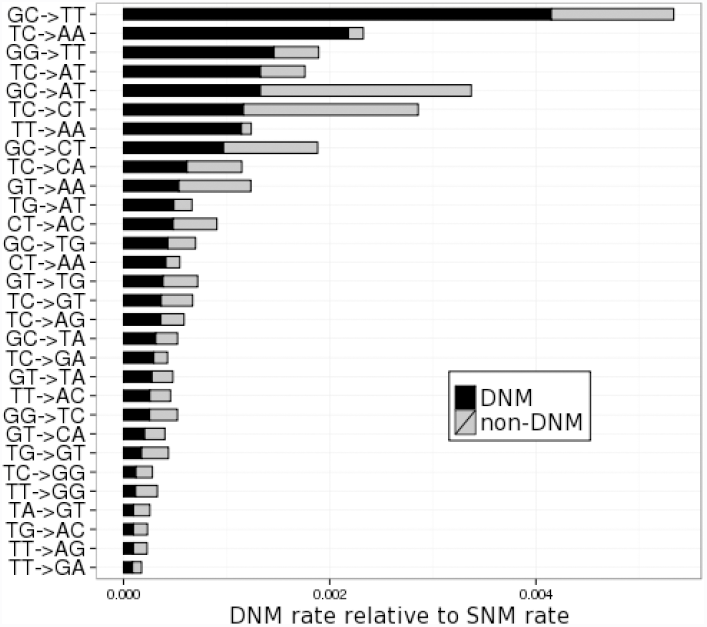
Top 30 most frequent DNMs. Reverse complementary DNMs were combined. DNM rate (black) varies between types of DNMs; GC->TT/AA DNM rate by far exceeds other rates.

### Pol ζ acts in late replicating regions

RT and replication by pol *ζ* are two factors affecting the accumulation of mutations. We investigated the relation between these factors by tracing the mutational signature of pol *ζ* along the genome. Unlike the overall DNM rate which is only weakly correlated with RT (Fig. 1D), the rate of GC->AA/TT DNMs is 61% higher in late RT (Fig 3B, p<2 *10^−6^, permuted CA test; Suppl. Table 2). Conceivably, this association could arise due to other genomic features correlated with RT. To test whether RT has an independent effect on the GC->AA/TT DNM rate, we analyzed it together with a set of other features that are correlated with the human mutation rate: GC content, H3K9me3 histone modification, density of DNase I hypersensitivity sites (DHS), and recombination rates (1000 Genomes Project Consortium 2010). We applied a Poisson regression model, using these values as well as the SNM rates and overall DNM rate as the explanatory variables, the number of GC->AA/TT mutations as the response variable, and the number of GC dinucleotides as the exposure. The contribution of RT remained highly significant (Suppl. Table 3, p < 3*10^−5^), indicating that RT is indeed independently associated with the mutational signature of pol *ζ*. Additionally, the genomic locations of GC->AA/TT TSBMs are biased towards late RT, compared to the GC->AA/TT TDBMs (p<0.01, Mann–Whitney U test).

To single out the signature of pol *ζ* from the overall rate of DNMs in relation to RT, we consider the ratio of the rates of GC->AA/TT DNMs and all DNMs. This ratio increases monotonically from early to late RT, differing between the regions of earliest and latest replication by 46% (Fig. 3C, p<6 *10^−4^, permutation test), further arguing for the distinctness of mutational mechanisms that underlie the signature of pol *ζ* and other DNMs.

**Figure 3.**
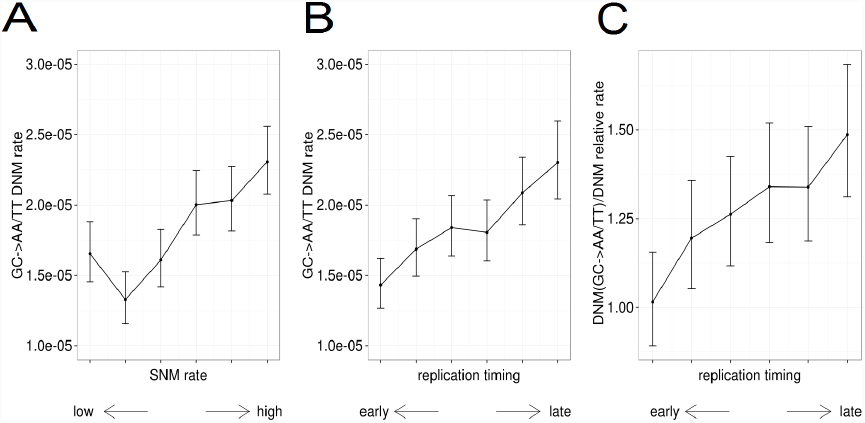
Association of GC->AA/TT DNM with replication timing and mutation rate. (A) GC->AA/TT DNM rate increases with SNM rate. (B) GC->AA/TT DNM rate increases with RT. (C) Ratio of GC-AA/TT DNM rate to DNM rate increases with RT.

Leading and lagging DNA strand in humans are synthesized by pol ε and pol δ respectively (Shinbrot et al. 2014). Distinct biological processes operating on these two stand types lead to differences in mutational footprints between the leading and the lagging strands (Chen et al. 2011; Baker et al. 2012). Since the signature of pol *ζ* is associated with RT, we asked whether it preferentially occurs on one type of strand. We use the derivative of RT with respect to the genomic coordinate as a proxy for the propensity of nucleotide to be replicated as the leading strand (Baker et al. 2012). We see no stand asymmetry with respect to the ratio of GC->AA and GC->TT mutations (Suppl. Fig S3); thus, the action of pol *ζ* is likely uncoupled from strand-specific replication mechanisms.

### Pol ζ signature is associated with transcription in a strand-asymmetric way

In yeast, pol ζ is involved in transcription-associated mutagenesis (TAM) (Datta and Jinks-Robertson 1995; Kim et al. 2007). Divergence is positively correlated with gene expression due to susceptibility of single stranded DNA to mutations during transcription (Park et al. 2012). The non-template strand is more frequently found in single stranded state during transcription than the template strand, leading to asymmetry of mutation patterns between strands (Polak and Arndt 2008). We observe that the GC->TT DNMs are 40% more frequent, compared to the GC->AA DNMs on the non-template strand, as well as to both types of mutations in intergenic regions (Fig. 4). This pattern further supports association of the GC->AA/TT DNMs with transcription, and implies that the GC->TT DNM rate is increased in transcribed regions of the non-template strand. The association of mutation rate with transcription may be direct, or may arise due to higher chromatin accessibility in the transcribed regions. However, we observe a negative correlation between the GC->AA/TT DNM rate and DNA accessibility estimated from the density of DNase I hypersensitivity sites (Thurman et al. 2012) (Suppl. Fig. 2). Therefore, pol ζ is probably recruited to damage-prone single-stranded DNA on the non-template strand, thus increasing the mutation rate during transcription.

**Figure 4.**
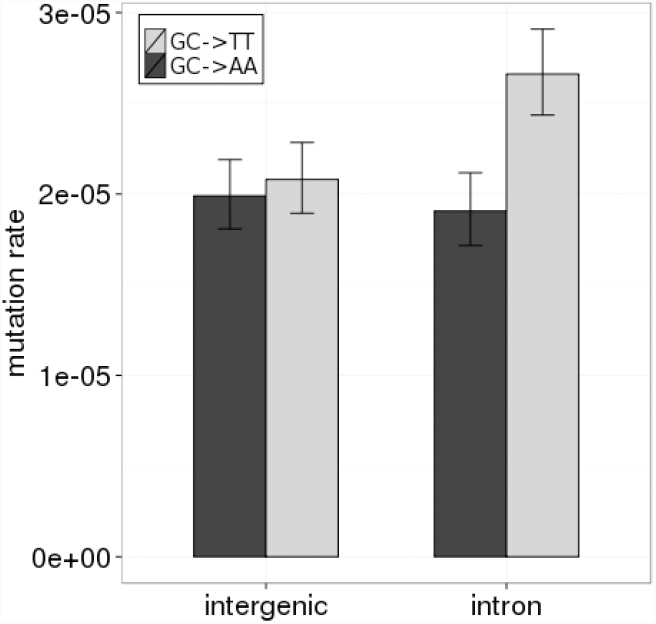
Strand-specific asymmetry of GC->AA/ TT DNMs in transcribed regions. The GC->TT DNMs rate significantly exceeds the complementary GC->AA DNMs rate in introns on the non-template strand. For intergenic regions, the direction of mutation is determined using the reference orientation.

### Spontaneous deamination recruits pol ζ

CpG dinucleotide is highly mutagenic in mammals due to spontaneous deamination of methylated cytosine (Bird 1980). Spontaneous deamination converts 5-methylcytosine to thymine, causing T-G mispairing. Mispaired thymine residues are then resected by thymine-DNA glycosylase (TDG) producing abasic sites (Gallinari and Jiricny 1996). Pol ζ plays an important role in abasic sites bypassing during replication in human cells (Weerasooriya et al. 2014). Following Giannoulatou et al. (2013), we asked whether the GC->AA/TT DNM rate increases if one of the nucleotides is in the CpG context. The CGC->CAA and GCG ->TTG DNMs are 29% more frequent than the GC->AA/TT DNMs out of the CpG context (p<0.05, permutation test). This effect could not be attributed to the elevated mutation rate in CpG, since estimation of DNM events accounts for CpG hypermutability through subtraction of TDBMs (see Methods). Therefore, pol ζ is actively involved in TLS on the strand complementary to the mutated CpG dinucleotides.

### Pol ζ is likely involved in the Costello syndrome

As one prominent example, we noticed that the DNM causing the Costello syndrome is GCG ->TTG (Giannoulatou et al. 2013) and therefore could be linked to pol ζ activity. GC ->TT DNM occurs in an exon of the HRAS gene, and causes the Gly12->Val12 substitution (Fig. 5B). This DNM is a frequent germline mutation; in fact, it is 1.7 times more frequent than a single nucleotide G35 ->T35 mutation which causes the same amino acid substitution (Giannoulatou et al. 2013). This means that the high prevalence of the GC ->TT DNM in HRAS cannot be explained by germline selection favoring the Gly12 ->Val12 substitution, and suggests that this DNM may arise through a non-canonical mutational mechanism. Pol ζ is an error-prone and transversion-prone (Zhong et al. 2006; Sakamoto et al. 2007; Stone et al. 2012) poorly processive polymerase (Nelson et al. 1996; Lee et al. 2014); thus, its recurrent activity in the first exon of the HRAS gene should cause a strong increase in the mutation rate and decrease in the transition/transversion ratio spanning a few hundred nucleotides nearby. Using the human-mouse divergence data, we estimated the local mutation rate near this site. In the two introns adjacent to the exon containing the site of the GC->TT DNM, the human-mouse divergence is increased by a factor of ~2, compared to the intron 1Kb away, where it is close to the chromosome average (Fig. 5A). Moreover, the fraction of transversions is also increased by a factor of 2, which is much higher than the 20% increment in usual mutational hotspots (Seplyarskiy et al. 2012). A twofold increase in divergence in two adjacent introns is also observed in the human-marmoset comparison (Suppl. Fig. 4). As the DNM is not fixed between species, the local increase in divergence is not a consequence of a past complex MNM involving the GC ->TT DNM. Bursts of substitutions in adjacent introns are hardly due to selection, because there are no amino acid substitutions in this exon, and the introns are not particularly conserved. Therefore, there exists a local mutational hotspot with drastically decreased transition/transversion ratio around the site of the recurrent germline GC->TT DNM, consistent with the expected signature of pol ζ. Possibly, pol ζ is recruited for TLS of the strand complementary to the abasic site when this site arises due to C_met_36->T36 deamination and subsequent T resection via TDG. This mechanism should lead to an excess of GC->TT over GC->AA, which is clearly observed in the Costello syndrome (Giannoulatou et al. 2013), although among point mutations, G->A is more frequent than G->T at HRAS G35 nucleotide (Giannoulatou et al. 2013).

**Figure 5.**
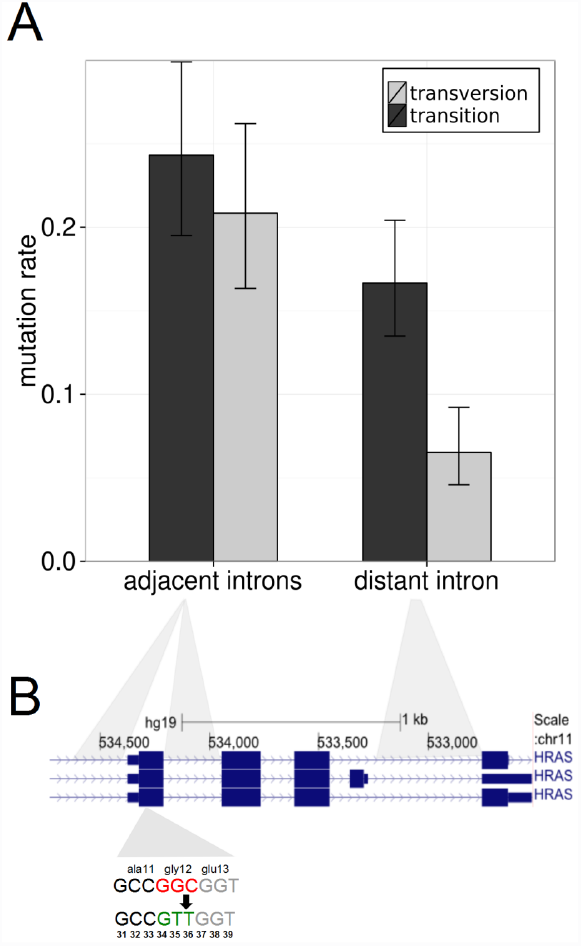
GC->AA/TT DNM in HRAS gene is a germline hotspot mutation associated with Costello syndrome. (A) Introns adjacent to the GC->TT DNM hotspot are substantially more divergent, and carry a larger fraction of transversions, compared to a more distant intron. (B) Structure of the HRAS transcript. The GC->TT DNM occurs in gly12 of the first exon.

### DNMs are associated with local mutation hotspots

The association of the recurrent GC->AA/TT DNM with a mutation hotspot in the HRAS gene prompted us to study the local mutational landscape of DNMs. To this end, we studied the rate of SNMs in the vicinity of TSBMs or TDBMs. The SNM rate is elevated in the vicinity of both TSBMs and TDBMs at distances of up to several kilobases (Fig. 6A). The increase in the vicinity of TSBMs could be due to association of DNMs with local mutational hotspots; however, the increase in the vicinity of both TSBMs and TDBMs could also be due to two co-occurring SNMs being markers of such hotspots (Johnson and Hellmann 2011; Seplyarskiy et al. 2012). We can infer the effect of DNMs per se from the known mutation rates in the vicinity of TDBMs and TSBMs and the fraction of DNMs among TSBMs (see Methods).

**Figure 6.**
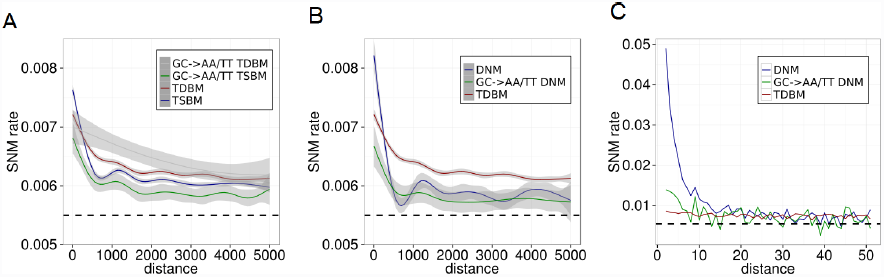
Local regions of DNMs are associated with mutation hotspots. The black dashed line is the genome average SNM rate. (A) The average SNM rate decreases with distance (2-5000 nucleotides) from tandem mutations (TDBM, TSBM, GC->AA/TT TDBM, GC->AA/TT TSBM). (B) The average SNM rate decreases with distance from DNMs. In (A) and (B), SNM rate was fitted by generalized additive linear model using cubic spline method. (C) Local decrease of SNM rate.

Somewhat unexpectedly, SNMs are less frequent around TSBMs than around TDBMs, except for the 10 nearest nucleotides (Fig.6A). Thus, contrary to naïve expectations, the occurrence of two mutations at adjacent sites in the human lineage is a poorer predictor of an SNM hotspot than the occurrence of two mutations at adjacent sites in different lineages. Still, the SNM rate is also increased in the vicinity of DNMs. The increase associated with the GC->AA/TT DNMs is rather similar to that associated with other DNM types (Fig. 6B).

The situation is somewhat different in the 10 nucleotides immediately adjacent to the TSBM or TDBM site (Fig.6C). At this distance, the rate of SNMs is radically increased around DNMs, but not around TDBMs; therefore, this increase is lineage-specific. This increase is likely due to a high fraction of complex mutations involving these DNMs, *e.g.* trinucleotide mutations (Terekhanova et al. 2013). Such complex mutations usually span distances of up to 20 nucleotides (Terekhanova et al. 2013), and are associated with a decreased transition/transversion ratio (Harris and Nielson 2014); both these features are observed in our data (Fig. 6C and Suppl. Fig. 5, respectively). These complex mutations are hardly produced by non-allelic gene conversion, since we exclude paralogous segments from analysis.

In contrast to DNMs overall, we observe only a very slight increase of the SNM rate near the GC->AA/TT DNM (Fig. 6C). This further supports the conjecture that this DNM usually occurs as the mutational signature of pol ζ, rather than via a mechanism common to the other DNMs and, in particular, as a part of more complex mutations.

In summary, the mutation rate in the vicinity of pol ζ signature is similar to that around other DNMs. This is in contrast to the pattern observed in the HRAS gene, where the DNM suggestive of pol ζ TLS occurs recurrently. These observations can be reconciled if most of the pol ζ-associated DNMs occur at random locations along the genome with only weak locational preferences, whereas a small fraction occurs at specific sites such as the HRAS gene over and over again, radically increasing the SNM rate in the vicinity.

## Discussion

Our analysis is based on overrepresentation of substitutions co-occurring in the same phylogenetic lineage. Besides simultaneous mutations, clustering of substitutions at one phylogenetic branch (Bazykin et al. 2004) or clustering of mutations at one haplotype (Callahan et al. 2011) may arise due to epistatic selection. Such selection-caused clustering, however, is expected to be the strongest in regions of the highest conservation (Halligan et al. 2011; Bazykin and Kondrashov 2012). By excluding exons, UTRs, and intron flanks, we do not consider most regions under strong selection. Other selectively important regions, which comprise up to ~8% of the human genome (Rands et al. 2014), are unlikely to affect our results. Non-allelic gene conversion may cause DNMs if the source sequence differs from the converted sequence in two tandem nucleotides. However, exclusion of 100 nucleotide regions centered at tandem mutations that have regions of high homology elsewhere in the human genome leads to only marginal differences in the GC->AA/TT signature and overall rate of tandem substitutions (Suppl. Fig. S5, S6), similar to our previous results (Terekhanova et al. 2013). As we do not consider sites masked by RepeatMasker, the observed trends are not due to mutational properties of repeats. In summary, our results are not an artifact of selection, gene conversion or repeats mutability.

In line with human diversity and disease studies (Chen et al. 2013; Harris and Nielsen 2014), GC->AA/TT DNM is the most frequent DNM in the human-chimp divergence. Simultaneous GC->AA/TT DNMs comprise the majority (79%) of observed TSBMs, and their rate exceeds the average DNM rate by a factor of ~10. Studying the genomic distribution and strand bias of these DNMs reveals a number of patterns. We found that the pol ζ mutational signature is strongly biased towards late replicating regions. This observation is independent of other factors driving variation of the human mutation rate (GC content, DNA accessibility, H3K9me3, recombination) as well as from variation of the SNM rate and DNM rate. The observed GC->AA/TT DNMs enrichment could be a result of decreased DNA repair fidelity, e.g. mismatch repair (Lujan et al. 2014), rather than reinforced recruitment of pol ζ. However, while the overall DNM rate could depend on DNA repair efficiency, there are no reasons to expect pronounced differences between the repair mechanisms leading to GC->AA/TT and all other DNMs.

Therefore, the most parsimonious explanation for higher GC->AA/TT DNM rate after normalizing by the overall DNM rate seems to be reinforced recruitment of pol ζ in late replicating regions. This implies that higher activity of error-prone polymerases is one of the causes of higher mutation rate in late RT. These results should not, however, be taken to imply that higher pol ζ activity in late RT is the main determinant of the relationship between RT and the mutation rate; more data is needed to estimate the fraction of genome replicated by pol ζ, and thus, the fraction of mutations resulting from pol ζ activity.

The error rate of pol ζ is several orders of magnitude higher than that of the major polymerases δ and ε (Zhong et al. 2006; Sakamoto et al. 2007). Thus, even a small fraction of DNA replicated by pol ζ may affect the genome average mutation rate. Heterogeneous recruitment of pol ζ affects the mutation rate along the genome. Local regions carrying the GC->AA/TT DNMs exhibit elevated mutation rates; however, this increase is slight. In contrast, in the example of HRAS were pol ζ is apparently recruited repeatedly, we see a nearly twofold increase in the mutation rate around the DNM. Therefore, most of the pol ζ mutations genomewide probably occur at more or less random locations, and are not markers of mutational hotspots.

The process of mutagenesis is linked to transcription. Several important factors underlie the mutation patterns of transcribed regions, including two conflicting ones: elevated mutagenesis of the non-template strand which is often in a single stranded state (Polak et al. 2008) and transcription-coupled repair of the template strand (*e.g.* Marteijn et al. 2014). The former factor predominates in the germline, leading to accelerated divergence of highly expressed genes (Park et al. 2012), while the latter predominates in cancer, leading to decreased density of somatic mutations in transcribed regions (Polak et al. 2014). Transcription may lead to strand-specific mutagenesis; the most prominent example is the asymmetry of A->G/T->C mutations, which is observed in the divergence of primates (Polak et al 2008). We observe GC->AA/GC->TT asymmetry in introns (Fig. 4), consistent with pol ζ-dependent mutagenesis in transcribed regions in humans, in line with previous findings in yeast (Datta and Jinks-Robertson 1995; Kim et al. 2007). Assuming that pol ζ is associated with the single-stranded non-template strand, we suggest that the true trace of pol ζ TLS is the GC->TT DNM.

The recurrent GC->TT DNM causing the Costello syndrome provides an opportunity to investigate a likely GC->TT hotspot. Besides the twofold increase in the mutation rate, we see a twofold excess of transversions over transitions at distances of up to 500 nucleotides, compared to more distant DNA segments. Both these patterns as well as the length of the regions that they span are in agreement with frequent pol ζ TLS of this region, pointing to pol ζ as a likely source of the mutation causing the Costello syndrome.

The GC->AA/TT DNMs have properties distinct from those of other DNMs. They show a much more radical association with replication time. While both GC->AA/TT DNMs and other DNMs reside in regions with elevated large-scale mutation rate as well as in local mutation hotspots, sites located within 10 nucleotides of non-GC->AA/TT DNMs tend to be surrounded by other mutations, while GC->AA/TT DNMs do not. This contrast is likely due to the fact that a large subset of non-GC->AA/TT DNMs (but not GC->AA/TT DNMs) arises from complex events comprising DNMs. This again distinguishes the GC->AA/TT DNM among other types of DNMs, in line with their origin from a distinct molecular mechanism.

In divergence of closely related species, multiple substitutions occurring at the same lineage at several adjacent sites represent complex events rather than sets of SNMs. Using genome-scale divergence data to infer the rates and types of complex mutations can be informative of the mechanism by which they originate. Reciprocally, combining the knowledge of specific processes that give rise to complex mutations of a specific kind with their genomic distribution is informative of the properties of the underlying mutational mechanism. Application of this approach to GC->AA/ TT DNM reveals the complexity of the genomic activity of pol ζ.

## Materials and methods

### Substitution data

To analyze mutational processes operating in the human genome, we identify putatively neutral sites. Multiple alignment of human, chimpanzee and orangutan genomes was downloaded from the UCSC genome browser (Karolchik et al. 2014). We exclude all exons included in the USCS knownGene annotation together with their 50 nucleotide flanks, 5’ and 3’ UTRs, nucleotide sites masked by RepeatMasker, and non-A, C, G or T nucleotides in any of the three genomes, as well as the X and Y chromosomes. We reconstruct the state of the human-chimpanzee common ancestor and map substitutions to either the human of the chimpanzee lineage using maximum parsimony (Fig 1A). Only sites with no more than one difference between human, chimpanzee and orangutan genomes were considered. In the main analyses, we exclude sites that overlap a CpG dinucleotide in the human-chimp common ancestor. In the analysis of multinucleotide mutations, we exclude tandem substitutions that may occur through the CpG intermediate state; *i.e.* we do not consider situations when in the first site, either the derived or the ancestral variant is ‘C’, and in the second site, either the derived or the ancestral variant is ‘G’. The point mutation rate of a genomic region was estimated as the number of substitutions divided by the number of sites.

### Dinucleotide mutations

Tandem same branch mutation (TSBM) was defined as two adjacent substitutions on the human branch, whereas tandem different branch mutation (TDBM) was defined as two adjacent substitutions such that the first occurred on the chimpanzee branch, and the second, on the human branch (Fig. 1A). The number of dinucleotide mutations (DNMs), *i.e.*, simultaneously occurring adjacent mutations, was estimated as the excess of TSBMs over TDBMs:

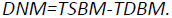

DNM and SNM rates are calculated as the number of DNM or SNM events divided by the number of considered dinucleotide sites or nucleotide sites respectively. To calculate this rate for each dinucleotide, we divided the number of considered DNMs by the appropriate number of ancestral dinucleotides. The GC->AA/TT DNM rate in the CpG context was determined as follows:

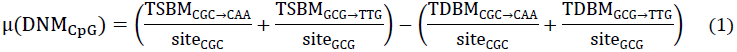

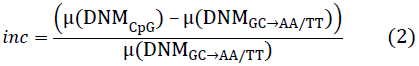

where *μ*(*DNM*_*CpG*_) is the DNM_GC->*AA/TT*_ rate in the CpG context, *site*_*CGC*_ and *site*_*GCG*_ are the numbers of corresponding trinucleotides, and *inc* is the relative increment of the DNM_GC->AA/TT_ rate in the CpG context over the DNM_GC->AA/TT_ rate.

To estimate the significance of the observed *inc* values, we used a bootstrapping procedure, sampling with replacement from the set of TSBMs from appropriate sites and calculating *inc* in 1000 bootstrap trials.

### Estimation of the mutation rate and transition/transversion ratio in HRAS gene

To analyze the local mutational hotspot in HRAS gene, we use human-mouse comparison. We compared the divergence rates in the two adjacent introns (hg19 coordinates: Chr11:534386-534585 and Chr11:534001-534200) and in the distant intron (Chr11:532941-533440).

### Local mutation patterns

We analyzed the SNM rate for nucleotide sites at 2-5000 nucleotides from the site of tandem substitution. DNMs that were involved in a trinucleotide mutation spanning three adjacent nucleotides were excluded. To infer the SNM rate around DNMs, we used the fraction of DNMs among TSBMs, and the SNM rate around TSBMSs and TDBMs:

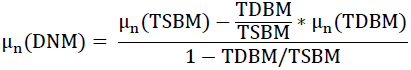

where μ_n_(DNM) is the local SNM rate at distance n from DNM, μ_n_(TSBM) is the local SNM rate at distance n from TSBM, and μ_n_(TDBM) is the local SNM rate at distance n from TDBM. Using similar equations, we obtained the SNM rate around GC->AA/TT DNMs, as well as the transition/transversion ratio around DNMs and GC->AA/TT DNMs.

### Genomic features

Replication timing was obtained from Koren et al. (2012). DNase I hypersensitivity sites and H3K9me3 of Gm12878 cell line were downloaded from ENCODE (https://www.encodeproject.org/). Recombination rates were obtained from (1000 Genomes Project Consortium 2010). All tracks were remapped to the human genome build of hg19, which was used in all analyses. Replication timing, DNase I hypersensitivity sites, H3K9me3, recombination rates, GC content, and SNM rates were averaged in 50 kb non-overlapping regions. We classified these regions into equal-sized bins according to the value of each genomic feature. Each analysis was performed using 3, 6, 8 and 10 bins to test the robustness of the results to the specific bin boundaries used. We then measured the DNM rate and GC->AA/TT DNM rate for each bin. To estimate the significance of the observed trends, we calculate the p-value of the Cochran-Armitage statistic for trend in proportions using the null distribution obtained from permutations. We randomly permute values of RT among 50 kb non-overlapping regions to obtain the permutation-based null distribution. To test whether the GC content acts as a confounder for association with replication timing, we preserved the GC distribution of each bin during permutation.

Null distribution was further fitted to Gamma distribution utilizing fitdistrplus R package, using the Kolmogorov-Smirnov test (K-S test) to estimate the adequacy of this procedure (K-S test resulted in p > 0.05 in all cases). The Q-Q plots showed that the p-values obtained with this procedure are conservative. Significance of the trend for the ratio of GC->AA/TT DNM rate and overall DNM rate was estimated from the distribution of the increments of this ratio from each bin to the next, obtained from 500 permutations.

For the regression analysis, for each 50 kb region, we collected a vector of averaged genomic features (replication timing, GC content, H3K9me3, DNase I hypersensitivity sites, recombination rate, SNM rate, DNM rate). These vectors were used as the input for Poisson regression, with the number of GC->AA/TT events as the response variable, and the number of GC dinucleotides as exposure. Poisson regression was performed using the R function glm().

## Acknowledgements

### Acknowledgments

We thank Andrey Mironov, Igor Rogozin, and Shamil Sunyaev for useful discussions, Nadezhda Terekhanova for data preprocessing, and Mikhail Gelfand for useful comments on the manuscript.

